# The WD40-repeat protein WDR-48 promotes the stability of the deubiquitinating enzyme USP-46 by inhibiting its ubiquitination and degradation

**DOI:** 10.1101/679415

**Authors:** Molly Hodul, Rakesh Ganji, Caroline L Dahlberg, Malavika Raman, Peter Juo

## Abstract

Ubiquitination is a reversible post-translational modification that has emerged as a critical regulator of synapse development and function. However, mechanisms that regulate the deubiquitinating enzymes (DUBs) that are responsible for the removal of ubiquitin from target proteins are poorly understood. We previously showed that the DUB USP-46 removes ubiquitin from the glutamate receptor GLR-1 and regulates it trafficking and degradation in *C. elegans*. We found that WD40-repeat proteins WDR-20 and WDR-48 bind and stimulate the catalytic activity of USP-46. Here, we identify another mechanism by which WDR-48 regulates USP-46. We found that increased expression of WDR-48, but not WDR-20, promotes USP-46 abundance in mammalian cells in culture and in *C. elegans* neurons *in vivo*. Inhibition of the proteasome promotes the abundance of USP-46, and this effect is non-additive with increased expression of WDR-48. We found that USP-46 is ubiquitinated, and expression of WDR-48 reduces the levels of ubiquitin-USP-46 conjugates and increases the half-life of USP-46. A point mutant version of WDR-48 that disrupts binding to USP-46 is unable to promote USP-46 abundance *in vivo*. Together, these data support a model in which WDR-48 binds and stabilizes USP-46 protein levels by preventing the ubiquitination and degradation of USP-46 in the proteasome. Given that a large number of USPs interact with WDR proteins, we propose that stabilization of DUBs by their interacting WDR proteins may be a conserved and widely used mechanism to control DUB availability and function.

---

Ubiquitination is a widely used post-translational modification that regulates a large variety of neuronal processes including synapse development and function (1–3). The covalent attachment of ubiquitin to lysine residues on substrates by ubiquitin E3 ligases has many consequences for target proteins including degradation in the proteasome, changes in protein trafficking, or altered function. Ubiquitination is a highly regulated and reversible process, where the removal of ubiquitin is achieved by a family of proteases called deubiquitinating enzymes (DUBs). Although there are about 600 ubiquitin ligases encoded by the human genome, there are only about 95 DUBs, suggesting that DUB function is tightly regulated (4,5). Indeed, increasing evidence shows that DUBs can be highly selective for substrates and appear to have very specific cellular functions (5–7). Additionally, because DUBs are intracellular proteases and the ubiquitination of proteins affects their function with profound cellular consequences, tight regulation of DUBs is needed to prevent indiscriminate proteolytic cleavage of ubiquitin conjugates (8).

The ubiquitin-specific protease (USP) family represents the largest sub-family of DUBs comprised of 56 members that regulate diverse cellular functions (4,7,9). Three related DUBs, USP-46, USP-12, and USP-1, have received particular attention due to their roles in regulating neuronal function, cell growth and division, and DNA damage (10,11). We previously showed that USP-46 regulates glutamate receptor levels in neurons to control glutamatergic behavior in *C. elegans* (12). USP-46 deubiquitinates the glutamate receptor GLR-1 and protects it from degradation in the lysosome. Similarly, in mammalian neurons, USP-46 can promote glutamate receptor subunit stability by deubiquitinating the glutamate receptor subunits GluA1 and GluA2 (13), indicating that this mechanism is conserved. Characterization of Usp46 mutant mice suggests that the DUB can also regulate GABA signaling and depression-like behaviors (14,15). In non-neuronal cells, USP-46 and USP-12 regulate a variety of cellular processes including cell proliferation and tumorigenesis (16–18). For example, USP-12 and USP-46 promote the stability of the Akt phosphatase PHLPP1 resulting in decreased cell proliferation and tumorigenesis in colon cancer cells (17,18). PHLPP1 is a tumor suppressor that has been implicated in several cancers including glioblastoma, colon and breast cancer (19–21). In contrast, overexpression of USP-12 promotes the stability and function of androgen receptors resulting in increased proliferation and survival of prostate cancer cells (16). Lastly, USP-1 is a critical mediator of two major DNA damage response pathways, the Fanconi anemia and DNA translesion synthesis pathways (22,23). The critical roles of these USPs in nervous system function, cell proliferation and cancer progression underscore the need to understand how these DUBs are regulated.

Recent work shows that DUBs can be regulated by a variety of mechanisms including transcription, post-translational modifications, and by interaction with other proteins (8,9). Large-scale proteomic studies revealed that the vast majority of DUBs interact with multiple proteins (24,25) that can regulate their subcellular localization, substrate recognition, and catalytic activity (8,9). Interestingly, 35% of USP enzymes interact with one class of protein, WD40-repeat (WDR) proteins, and 45% of those interact with multiple WDR proteins (10,24). The WDR domain forms a rigid β-propeller structure that provides a stable binding surface for protein-protein interactions (26,27).

In this study, we investigate the regulation of USP-46 protein levels by the WDR proteins WDR-48 (also known as USP-1-associated factor 1, UAF1) and WDR-20. USP-46 and its homolog USP-12 have low intrinsic catalytic activity (28–30). Biochemical studies showed that WDR-48 interacts with USP-46, USP-12 and USP-1 and stimulates their catalytic activity (24,28,30–32). WDR-20 forms a ternary complex with WDR-48 and USP-46/USP-12, but not USP-1 (24,29), and further enhances their catalytic activity *in vitro* (29,30). The mechanisms underlying the ability of the WDR proteins to activate USP-12 and USP-46 were recently revealed by their crystal structures. The crystal structures of both DUBs were solved in complex with WDR-48 (33–35), and USP-12 was additionally solved in a ternary complex with WDR-48 and WDR-20 (34). Together, these structural studies revealed that the WDR proteins interact at a site distal to the active site, suggesting an allosteric mechanism underlies the ability of the WDR proteins to stimulate catalytic activity of the DUBs.

Here we identify another mechanism by which WDR proteins regulate DUBs. We found that USP-46 is ubiquitinated and degraded in the proteasome. We show that WDR-48 promotes USP-46 protein abundance by binding to the DUB and inhibiting its ubiquitination. We propose that binding of WDR proteins to DUBs may provide a general mechanism to regulate the stability and thus availability of DUBs to carry out their various cellular functions.

## Results

### WDR-48 promotes USP-46 protein abundance in vivo

We previously showed that the *C. elegans* WD40-repeat proteins WDR-48 and WDR-20 bind to the deubiquitinating enzyme USP-46 and increase its catalytic activity (36). We noticed that expression of WDR-48 but not WDR-20 increases USP-46 protein levels in HEK293T cells. To test whether this effect of WDR-48 occurs in neurons *in vivo*, we generated transgenic worms (*pzIs40*) expressing GFP-tagged USP-46 (USP-46∷GFP) under control of the *nmr-1* promoter (37) in ventral cord interneurons where the WDR proteins and USP-46 are known to act (12,36). GFP-tagged USP-46 is functional because this transgene rescues specific glutamatergic behavioral defects observed in *usp-46* null mutants (Supplemental Fig. 1, see Experimental Procedures). We first tested whether expression of the WDR proteins in ventral cord interneurons altered total USP-46∷GFP levels by immunoblotting total worm lysates with anti-GFP antibodies. Consistent with our previous findings in HEK293T cells (36), we found that overexpression of WDR-48 alone (*wdr-48*(*xs*)) or WDR-48 and WDR-20 together (*wdr-48*(*xs*);*wdr-20*(*xs*)) in GLR-1-expressing interneurons increased USP-46∷GFP protein levels. In contrast, overexpression of WDR-20 alone (*wdr-20*(*xs*)) had no effect on USP-46∷GFP protein levels (Fig. 1, *A* and *B*). Next, we directly measured the levels of USP-46∷GFP fluorescence in *glr-1*-expressing neurons in the ventral nerve cord (VNC) of *C. elegans*. In wild-type animals, USP-46∷GFP is localized in a diffuse pattern throughout the cell bodies and ventral nerve cord processes (Fig. 1*C*). We estimated the levels of USP-46∷GFP protein in these neurons by measuring the average fluorescence of USP-46∷GFP in a defined anterior region of the VNC or in the cell body of the VNC neuron PVC (see Experimental Procedures). We found that overexpression of WDR-48 and WDR-20 in these interneurons increases USP-46∷GFP abundance by 2.6-fold (Fig. 1, *C* and *D*). This effect can be largely attributed to WDR-48, as overexpression of WDR-48 alone results in a 2.2-fold increase in USP-46∷GFP levels, whereas overexpression of WDR-20 alone results in a smaller increase of a 1.4-fold in USP-46∷GFP levels (Fig. 1, *C* and *D*). We observed similar effects in the soma of VNC neurons (Fig. 3B), suggesting that the WDR proteins increase USP-46∷GFP levels throughout the neuron.

**Figure 1.**
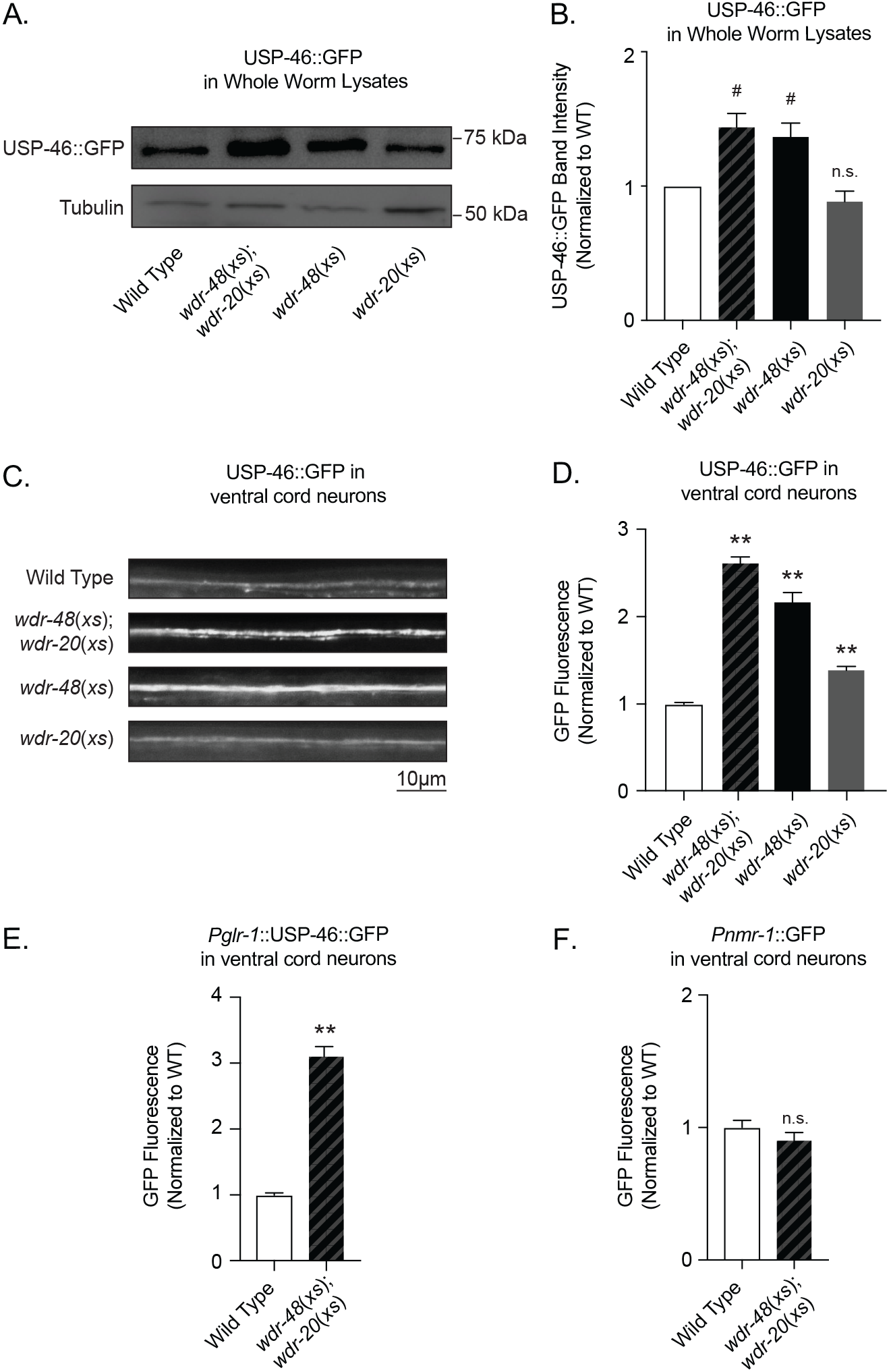
WDR-48 promotes USP-46 protein abundance. *A*, Representative immunoblots showing USP-46∷GFP expression levels (top), as detected with anti-GFP antibodies, in total worm lysates isolated from L4 stage larval animals harboring a USP-46∷GFP transgene expressed under control of the *nmr-1* promoter (*pzIs40*) in either a wild type background, or in animals overexpressing *wdr-48* and *wdr-20* together (*wdr-48*(*xs*)*;wdr-20*(*xs*)), *wdr-48* alone (*wdr-48*(*xs*)), or *wdr-20* (*wdr-20*(*xs*)) alone. Tubulin was also detected in these lysates (bottom) as a loading control. *B*, Quantification of the USP-46∷GFP bands from three independent western blots from the strains described in *A*. *C*, Representative images of USP-46∷GFP in the VNCs of L4 stage larval *pzIs40* animals for wild-type (n=57), *wdr-48*(*xs*)*;wdr-20*(*xs*) (n=49), *wdr-48*(*xs*) (n=16), and *wdr-20*(*xs*) (n=27) animals. *D*, Quantification of USP-46∷GFP fluorescence intensities (normalized) for the strains described in *C*. *E*, Quantification of the USP-46∷GFP fluorescence intensities in the VNCs of L4 larval animals harboring a USP-46∷GFP transgene expressed under the control of the *glr-1* promoter (*pzIs37*). Shown are the GFP fluorescence intensities (normalized) for wild-type (n=20) and *wdr-48*(*xs*)*;wdr-20*(*xs*) animals (n=20)*. F*, Quantification of VNCs of L4 larval animals harboring a GFP transgene expressed under the control of the *nmr-1* promoter (*pzEx386*). Shown are the GFP fluorescence intensities (normalized) for wild-type (n=47) and *wdr-48*(*xs*)*;wdr-20*(*xs*) animals (n=28). For all graphs, mean intensity ± S.E.M. are shown. Values that differ significantly from the wild type (Student’s *t*-test or ANOVA followed by Dunnett’s multiple comparison tests) are indicated as follows: #, p<0.05; **, *p* ≤ 0.001; n.s., p>0.05.

Because our USP-46∷GFP fluorescence reporter is expressed under control of the *nmr-1* promoter, the effects of the WDR proteins on USP-46 protein levels could be indirectly due to increased *nmr-1* promoter activity. We performed two experiments to test this possibility. First, we analyzed the effects of the WDR proteins on a second USP-46∷GFP reporter transgene (*pzIs37*) under control of an independent promoter, the *glr-1* promoter, which is also expressed in the ventral cord interneurons (38–40). We found that expression of WDR-20 and WDR-48 (*wdr-48*(*xs*);*wdr-20*(*xs*)) resulted in a similar 3-fold increase in USP-46∷GFP fluorescence in the VNC using this independent transgene (Fig. 1*E*). Second, we explicitly measured the activity of the *nmr-1* promoter using a transcriptional reporter (*Pnmr-1*∷GFP). We found that co-expression of WDR-48 and WDR-20 had no effect on GFP fluorescence (Fig. 1*F*). These data, together with the fact that we previously observed similar effects of the WDR proteins on USP-46 protein levels in HEK293T cells (where USP-46 was expressed using the mammalian expression vector pMT3)(36), suggest that the WDR proteins increase the abundance of USP-46 protein.

### USP-46 is regulated by the proteasome

We next sought to determine the mechanism by which WDR-20 and WDR-48 regulate USP-46 protein abundance by testing the hypothesis that the WDR proteins promote the stability of the DUB. We first measured the half-life of transiently transfected FLAG-tagged *C. elegans* USP-46 (FLAG-USP-46) in HEK293T cells. We blocked protein synthesis with the translational inhibitor cycloheximide (CHX) and measured the levels of FLAG-tagged USP-46 over time by western blotting with an anti-FLAG antibody. We found that USP-46 is relatively unstable and is degraded over time with a half-life of about 3-4 hours (Fig. 2, *A* and *B*, Fig. 3*A*). We next tested if USP-46 is degraded by the proteasome by co-incubating CHX-treated cells with the proteasome inhibitor bortezomib (BTZ). Compared to cells treated with CHX alone, we found that FLAG-tagged USP-46 was degraded more slowly over time and had an increased half-life (>8 hours) in the presence of CHX and BTZ (Fig. 2, *A* and *B*). To test whether USP-46 is also regulated by the proteasome in neurons *in vivo*, we treated USP-46∷GFP expressing worms with BTZ for 6 hours and measured USP-46∷GFP fluorescence in the soma of the VNC neuron PVC. We found that USP-46∷GFP levels increase about 3-fold in the presence of BTZ (Fig. 2*C*). Together, these data suggest that USP-46 protein stability is regulated by the proteasome.

**Figure 2.**
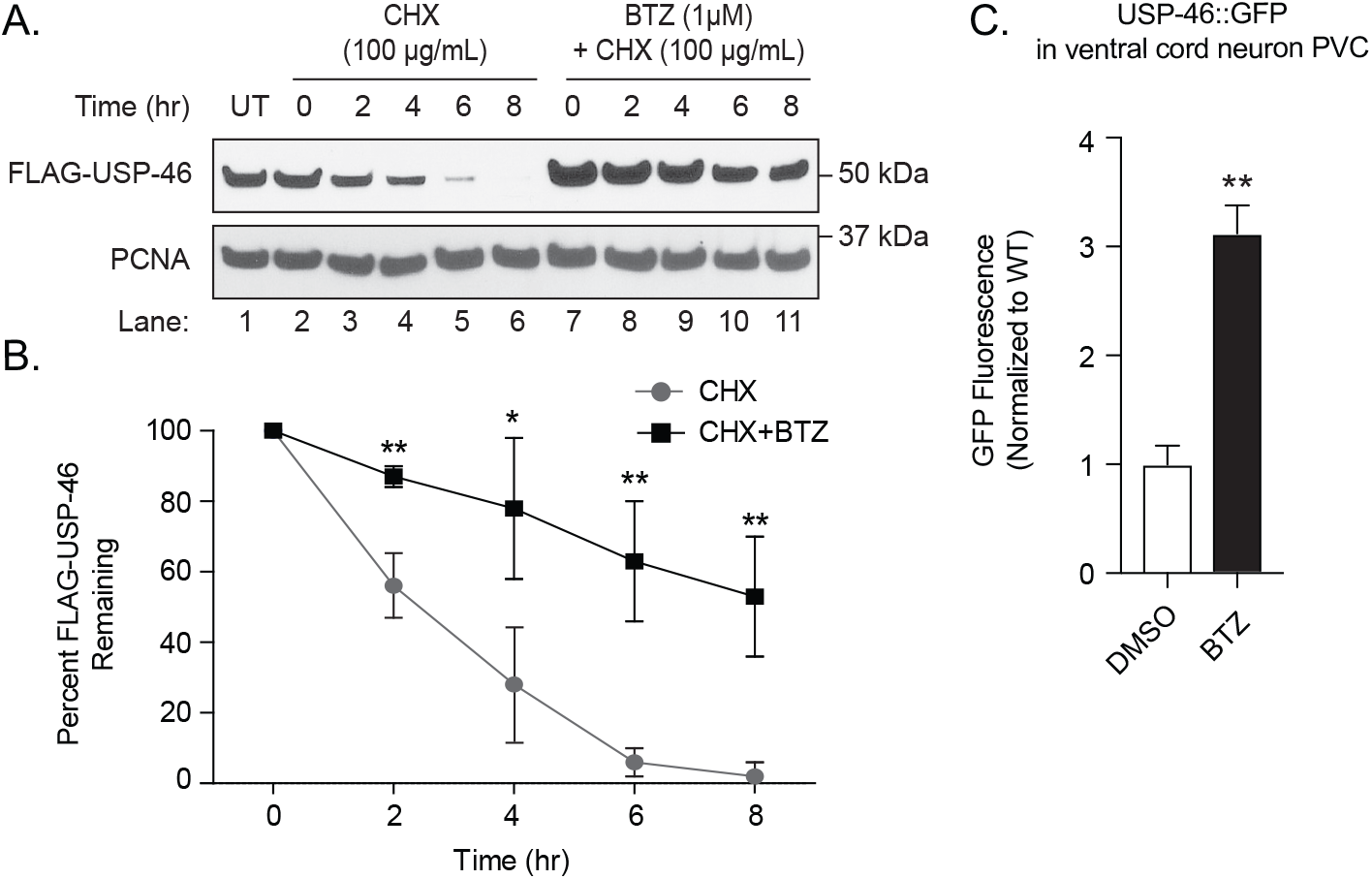
USP-46 is regulated by the proteasome. *A*, Representative immunoblot analysis of HEK293T cells transiently transfected with FLAG-USP-46 and either left untreated (UT, lane 1), or treated for the indicated times with cycloheximide (CHX, 100 μg/mL) alone (lanes 2-6) or CHX together with bortezomib (BTZ, 1 μM) (lanes 7-11). Cell lysates were immunoblotted for FLAG-USP-46 or PCNA (as a loading control). *B*, Quantification of the levels of FLAG-USP-46 from three independent experiments as described in A (normalized) are shown (means ± S.D.). *C*, Quantification of USP-46∷GFP fluorescence intensities (normalized) in the soma of PVC ventral cord neurons of L4 larval animals harboring a USP-46∷GFP transgene expressed under the control of the *nmr-1* promoter (*pzIs40*) treated with vehicle alone (DMSO) (n=12) or 50 μM BTZ (n=26, means ± S.E.M.) for 6 hours. Values that differ significantly from the control (Student’s *t*-test) are indicated as follows: *, *p* ≤ 0.01; **, *p* ≤ 0.001.

**Figure 3.**
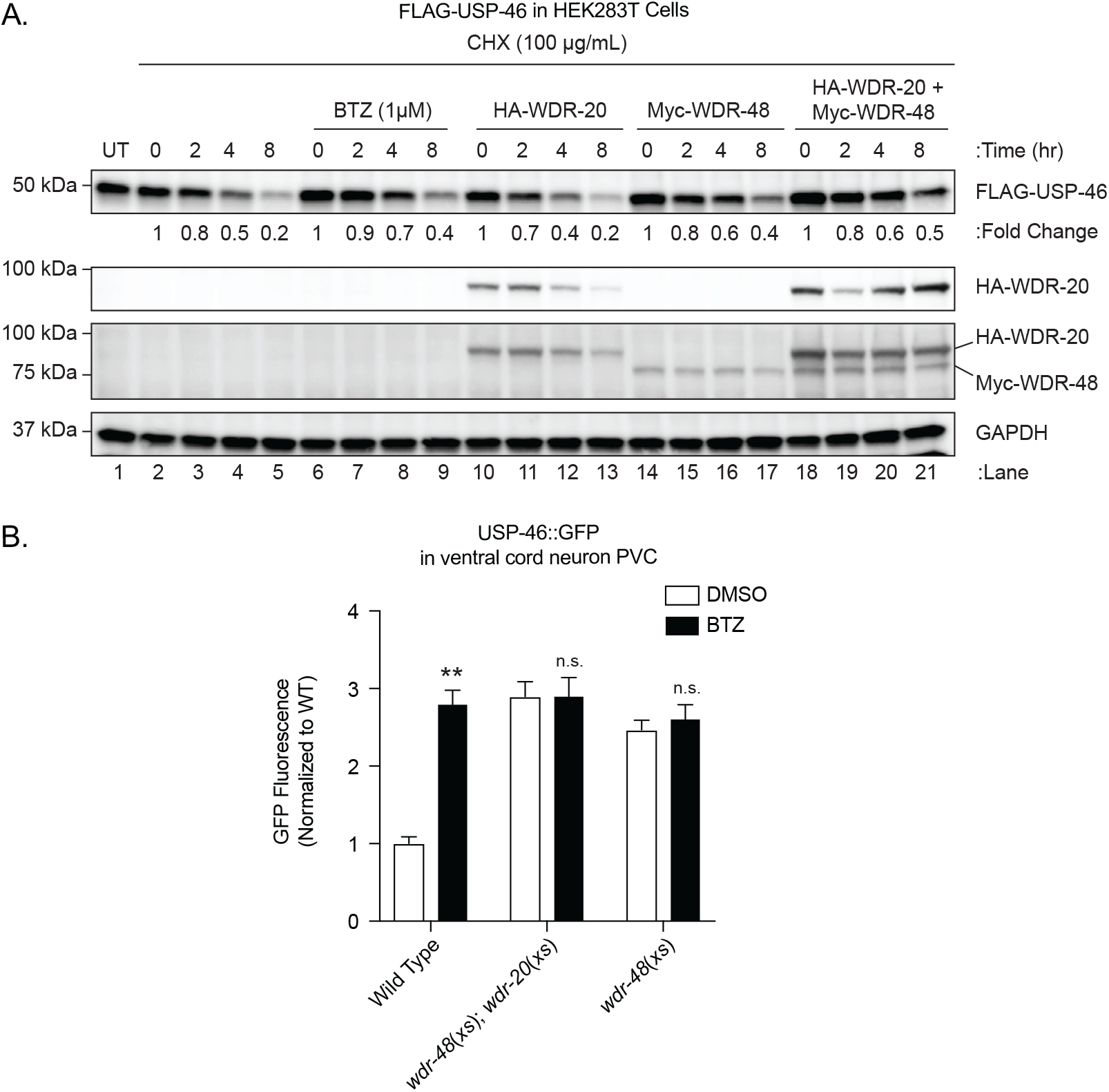
WDR-48 increases USP-46 protein stability. *A*, Representative immunoblot analysis of HEK-293T cells transiently transfected with FLAG-USP-46 and either left untreated (UT, lane 1) or treated with cycloheximide (CHX, 100 μg/mL) for the indicated times (lanes 2-21). In addition, cells were either treated with bortezomib (BTZ, 1 μM) (lanes 6-9) or transiently co-transfected with HA-WDR-20 (lanes 10-13), Myc-WDR-48 (lanes 14-17), or both HA-WDR-20 and Myc-WDR-48 together (lanes 18-21). GAPDH was used as a loading control. Whole cell lysates were immunoblotted for the various epitope-tagged proteins or GAPDH, as indicated. *B*, Quantification of USP-46∷GFP fluorescence intensities (normalized) in the soma of PVC ventral cord neurons of L4 larval animals harboring a USP-46∷GFP transgene expressed under the control of the *nmr-1* promoter (*pzIs40*) with or without 6 hour exposure to BTZ. Shown are the USP-46∷GFP fluorescence intensities for wild-type animals treated with either DMSO vehicle control (n=21) or BTZ (n=10), *wdr-48*(*xs*)*;wdr-20*(*xs*) animals in DMSO (n=10), *wdr-48*(*xs*);*wdr-20*(*xs*) animals in BTZ (n=8), *wdr-48(xs)* animals in DMSO (n=10), and *wdr-48*(*xs*) animals in BTZ (n=8). Both *wdr-48* and *wdr-20* were expressed under control of the *glr-1* promoter. BTZ treatment values that differ significantly from DMSO control within genotypes (ANOVA followed by Dunnett’s multiple comparison test) are indicated as follows: **, *p* ≤ 0.001; n.s., p>0.05.

### WDR-48 increases USP-46 protein stability

We next tested whether the WDR protein-mediated increase in USP-46 levels we observed (Fig. 1) is due to increased USP-46 stability. We measured levels of transiently transfected FLAG-tagged USP-46 over time in HEK293T cells treated with CHX in the presence or absence of HA-WDR-20 alone, Myc-WDR-48 alone, or HA-WDR-20 and Myc-WDR-48 together. Similar to the effect of BTZ on USP-46 protein levels (Fig. 3*A*, lanes 6-9), we found that expression of Myc-WDR-48 alone (lanes 14-17), or HA-WDR-20 together with Myc-WDR-48 (lanes 18-21), increased the stability of FLAG-USP-46 compared to CHX treatment alone (lanes 2-5). This effect is most evident at the 8 hour time point (compare lanes 5, 9, 17 and 21). In contrast, expression of HA-WDR-20 alone (lanes 10-13) did not stabilize FLAG-USP-46 resulting in a similar decline in USP-46 levels as observed with CHX treatment alone. Because USP-46 is degraded in the proteasome, we wanted to determine if the promotion of USP-46 abundance via WDR-48 is acting through the same mechanism *in vivo*. As described above, treatment of transgenic animals expressing USP-46∷GFP in VNC interneurons with BTZ results in a 3-fold increase in USP-46∷GFP fluorescence (Fig. 3*B*). Co-expression of WDR-48 alone (*wdr-48*(*xs*)), or WDR-48 and WDR-20 together (*wdr-48*(*xs*);*wdr-20*(*xs*)) result in similar increases in USP-46∷GFP fluorescence levels, and in the presence of the proteasome inhibitor BTZ, expression of WDR-48 does not confer any additional stability (Fig. 3*B*). Together, these data suggest that WDR-48 acts to reduce the proteasomal degradation of USP-46.

### WDR-48 inhibits ubiquitination of USP-46

If USP-46 is degraded by the proteasome, we would expect the DUB to be directly regulated by ubiquitin (Ub). We tested this idea using a HEK293T cell line stably expressing FLAG-tagged human USP-46. We treated these cells with BTZ for 6 hours to block degradation by the proteasome, immunoprecipitated FLAG-USP-46 under denaturing conditions, and used anti-Ubiquitin antibodies to probe for Ubiquitin-USP-46 conjugates. We found a low level of Ubiquitin-USP-46 conjugates in the absence of BTZ (Fig. 4*A*, lane 1), however treatment with BTZ results in readily detectable high molecular weight Ub-USP-46 conjugates (lane 2) consistent with polyubiquitination of USP-46. Interestingly, expression of Myc-WDR-48 alone (lane 4) or Myc-WDR-48 together with HA-WDR-20 (lane 5) in the presence of BTZ results in decreased levels of Ubiquitin-USP-46 conjugates compared to cells treated with BTZ alone (lane 2). In contrast, expression of HA-WDR-20 by itself (lane 3) did not reduce the levels of Ubiquitin-USP-46 conjugates, suggesting that WDR-48 specifically blocks ubiquitination of USP-46. Together, our data are consistent with the idea that USP-46 is ubiquitinated and degraded in the proteasome, and that WDR-48 inhibits USP-46 ubiquitination to promote its availability in the cell.

**Figure 4.**
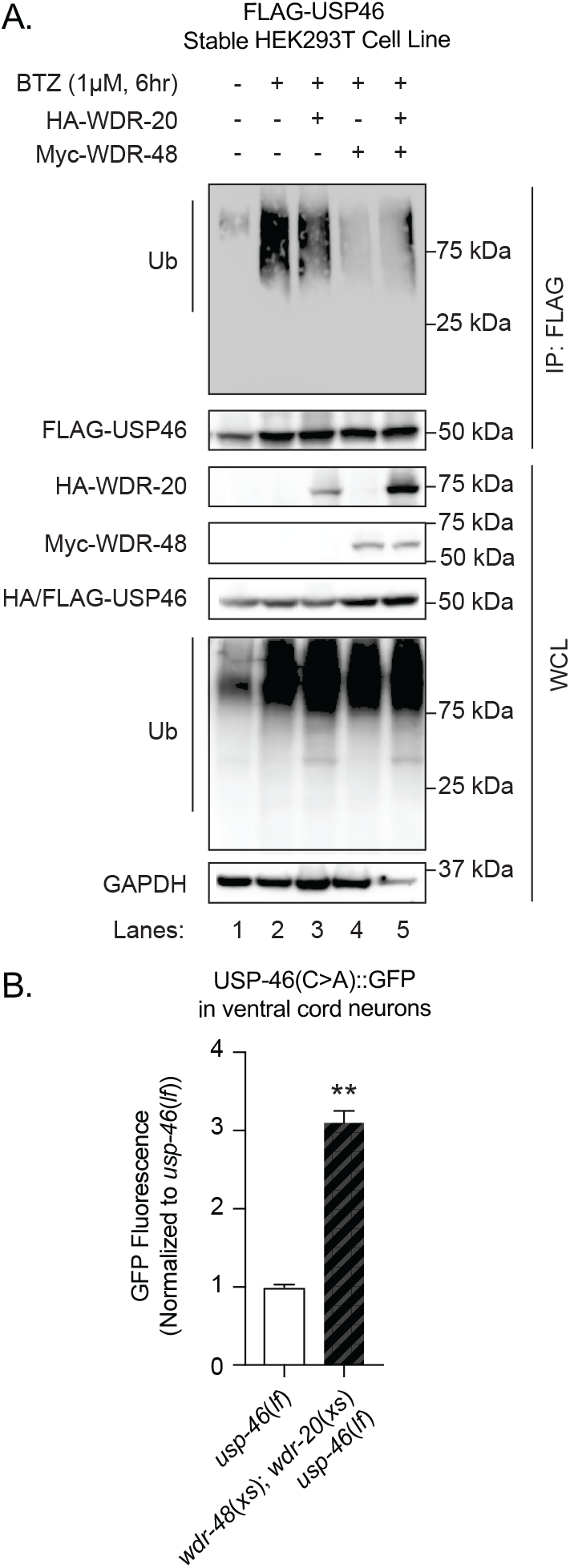
WDR-48 blocks ubiquitination of USP-46. Representative immunoblot comparing the levels of ubiquitin-USP-46 conjugates in HEK293T cells under various conditions. HEK293T cells stably expressing HA/FLAG-tagged human USP46 (HA/FLAG-USP46) were either left untreated (*lane 1*) or treated with Bortezomib (BTZ, 1 μM) for 6 hours (*lanes 2-5*). Some cells were also transiently transfected with either HA-WDR-20 (*lane 3*) alone, Myc-WDR-48 (*lane 4*) alone, or Myc-WDR-48 and HA-WDR-20 together (*lane 5*). Cell lysates were subjected to immunoprecipitation with anti-FLAG antibody under denaturing conditions followed by immunoblot analysis with anti-ubiquitin antibodies or anti-HA/FLAG antibodies, as indicated. Whole cell lysates (WCL) were immunoblotted for the various epitope-tagged proteins and GAPDH, as indicated. Similar results were obtained in three independent experiments. *B*, Quantification of GFP fluorescence intensity (means ± S.E.M.) in the VNCs of L4 larval *usp-46(ok2232)* null mutants harboring a USP-46(C38A)∷GFP transgene expressed under the control of the *glr-1* promoter (*pzEx378*) either without (n=19) or with co-expression of *wdr-48* and *wdr-20* (*wdr-48*(*xs*)*;wdr-20*(*xs*))(n=20). Values that differ significantly from the control (Student’s *t*-test) are indicated as follows: **, *p* ≤ 0.001.

We, and others, previously showed that WDR-48 and WDR-20 bind to USP-46 and increase its catalytic activity (28–30,36). Several USPs, such as USP4 and USP37, have been shown to auto-deubiquitinate *in trans* (41,42). Thus, we tested whether the ability of WDR-48 and WDR-20 to increase USP-46∷GFP levels *in vivo* was due to increased auto-deubiquitination of USP-46. We mutated the active site cysteine of USP-46, which is known to eliminate the catalytic activity of the DUB (36), and expressed catalytically-inactive USP-46(C>A)∷GFP in an *usp-46* null mutant background. We found that co-expression of WDR-48 and WDR-20 (*wdr-48*(*xs*);*wdr-20*(*xs*)) was still able to increase the abundance of USP-46(C>A)∷GFP in the VNC by 3-fold (Fig. 4*B*), which is similar in magnitude to the effect we observed on wild-type USP-46∷GFP (Fig. 1*D*). These results suggest that the WDR proteins increase USP-46 levels independent of its own catalytic activity.

### WDR-48 binding to USP-46 is required to stabilize the DUB

WDR-48, WDR-20 and USP-46 form a complex *in vitro* (24,28,29,36), and recently, the crystal structures of mammalian WDR-48/USP-46, WDR-48/USP-12 and WDR-48/WDR-20/USP-12 were recently solved (33–35). *C. elegans* USP-46 is the sole homolog of both mammalian USP-46 and USP-12, which share 90% sequence similarity with each other (12). The crystal structures defined the critical interaction interface between WDR-48 and USP-46/USP-12 revealing several key amino acid residues that are required for the interaction. Yin et al. showed that mutation of 3 key residues in WDR-48 (K214E, W256A, R272D) at the interface with USP-46 disrupted binding between the two proteins (35). These three residues are conserved in *C. elegans* WDR-48 as either identical or similar amino acids (Fig. 5*A*). We mutated these three residues in *C. elegans* WDR-48 (R224E, W266A, K282D, hereafter referred to as WDR-48(3Xmut)) to test if they would disrupt binding between WDR-48 and USP-46, as predicted by the high homology of these residues. We co-transfected HEK293T cells with FLAG-USP-46 and either wild-type Myc-WDR-48 or Myc-WDR-48(3Xmut). As we showed previously (36), FLAG-USP-46 co-immunoprecipitates with wild-type Myc-WDR-48 (Fig. 5*B*, lane 1). In contrast, a much lower amount of Myc-WDR-48(3Xmut) is pulled down with FLAG-USP-46 (Fig. 5*B*, lane 2), suggesting that the triple point mutant has a diminished ability to interact with USP-46. We next tested whether the ability of WDR-48 to promote USP-46 protein levels required a direct interaction between the two proteins. We generated transgenic animals expressing wild-type Myc-WDR-48 or Myc-WDR-48(3Xmut), under control of the *glr-1* promoter in the VNC neurons. Consistent with our previous data, we found that overexpression of wild-type Myc-WDR-48 results in increased levels of USP-46∷GFP as determined by western blotting of whole worm lysates (Fig. 5*C*) and by measuring USP-46∷GFP fluorescence in the VNC (Fig. 5*D*). In contrast, overexpression of Myc-WDR-48(3Xmut) was unable to increase levels of USP-46∷GFP (Fig. 5, *C* and *D*). The inability of Myc-WDR-48(3Xmut) to increase USP-46∷GFP levels *in vivo* is not due to differences in expression levels because the wild-type Myc-WDR-48 and Myc-WDR-48(3Xmut) transgenes were expressed at comparable levels as assessed by western blotting whole worm lysates (Fig. 5*E*). These data suggest that direct binding of WDR-48 to USP-46 is required for the ability of WDR-48 to promote USP-46 protein levels in neurons *in vivo*.

**Figure 5.**
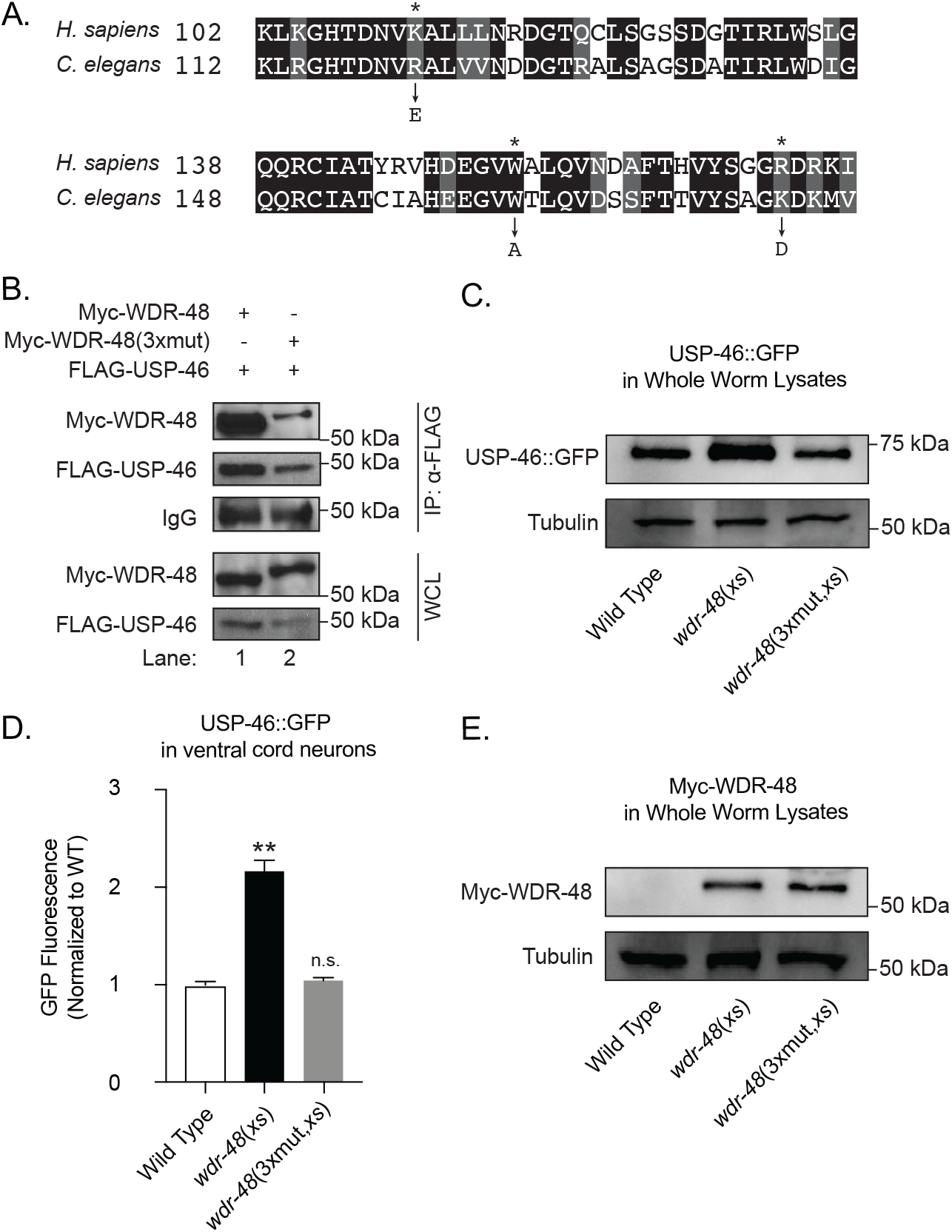
Direct binding of WDR-48 to USP-46 is required to stabilize the DUB. *A*, Partial protein sequence alignment illustrating the similarity (grey shading) or identity (black shading) of amino acids between *H. sapiens* and *C. elegans* WDR48 in the region that interacts with USP46 (34). Three point mutations in *H. sapiens* WDR48 (K214E, W256A, and R272D) that disrupt binding to the DUB are marked by arrows. The positions of the corresponding amino acids in *C. elegans* WDR-48 that were mutated in Myc-WDR-48(R224E, W266A, and K282D), also referred to as Myc-WDR-48(3Xmut), are marked by asterisks. *B*, Representative immunoblot showing the amount of Myc-WDR-48 and Myc-WDR-48(3Xmut) that was co-immunoprecipitated by FLAG-USP-46 from HEK293T cells. Cells were transiently co-transfected with FLAG-USP-46 together with either wild type Myc-WDR-48 (*lane 1*) or Myc-WDR-48(3Xmut, *lane 2*). Cell lysates were subjected to immunoprecipitation with anti-FLAG antibodies followed by immunoblotting with anti-Myc antibodies. Whole cell lysates (WCL) were immunoblotted for the various epitope-tagged proteins, as indicated. Similar results were obtained in three independent experiments. *C*, Representative immunoblot for total USP-46∷GFP, as detected with anti-GFP antibodies, in whole worm lysates of L4 larval animals harboring a USP-46∷GFP transgene expressed under the control of the *nmr-1* promoter (*pzIs40*, top) from wild-type animals, or animals expressing either wild type *wdr-48* (*xs*) or *wdr-48* (3xmut, xs) expressed under control of the *glr-1* promoter. Tubulin was also detected in these lysates (bottom) as a loading control. *D*, Quantification of USP-46∷GFP fluorescence in the VNCs of L4 larval animals harboring a USP-46∷GFP transgene expressed under the control of the *nmr-1* promoter (*pzIs40*). Shown are USP-46∷GFP fluorescence intensities (normalized) for wild type(n=15), wdr-48(xs, n=16) and *wdr-48*(3xmut, *xs*, n=12) (means ± S.E.M). *E*, Representative immunoblots for Myc-WDR-48 and Myc-WDR-48(3Xmut) expression, as detected with anti-Myc antibodies, of lysates from L4 larval animals of wild-type, *wdr-48(xs)*, and *wdr-48(3xmut,xs)* animals (top). Tubulin was also detected in these lysates (bottom) as a loading control. Results from three independent experiments show that the relative abundance of Myc-WDR-48 is similar to that of Myc-WDR-48(3xmut, means ± S.E.M.). Values that differ significantly from the wild type (ANOVA followed by Dunnett’s multiple comparison test) are indicated as follows: **, *p* ≤ 0.001; n.s., p>0.05.

## Discussion

Ubiquitin can be added and removed from target proteins, however less is known about the regulatory mechanisms that control deubiquitinating enzymes. We previously showed that USP-46 regulates glutamate receptor levels in the VNC of *C. elegans* and that two WDR proteins bind and stimulate the catalytic activity of USP-46 (12,36), consistent with other studies (24,28,29). Here, we identify another mechanism by which the WDR proteins regulate DUBs. We find that expression of WDR-48, but not WDR-20, increases the abundance of USP-46 in cultured mammalian cells and *C. elegans* neurons *in vivo*. We show that USP-46 is ubiquitinated and degraded in the proteasome. Overexpression of WDR-48 blocks ubiquitination of USP-46, and binding of WDR-48 to USP-46 is required to promote DUB protein levels. These data show that WDR-48 stabilizes USP-46 and suggest that controlling WDR-48 expression may provide a novel mechanism to control DUB availability.

What is the mechanism by which WDR-48 decreases ubiquitination of USP-46? Our data show that expression of WDR-48 decreases levels of ubiquitin-USP-46 conjugates (Fig. 4*A*) and that WDR-48 binding to USP-46 is required for its ability to stabilize the DUB (Fig. *5*). Because WDR proteins can stimulate the catalytic activity of USP-1, USP-12 and USP-46 (28,30,31,36) and several DUBs such as USP4 and USP37 have been shown to deubiquitinate themselves in *trans* (41,42), we tested if USP-46 catalytic activity was required for the ability of WDR-48 to regulate USP-46 levels. We found that expression of WDR-48 increased the levels of catalytically-inactive USP-46(C>A)∷GFP by about 3 fold (Fig. 4*B*), which is similar to the magnitude of its effect on wild-type USP-46∷GFP (Fig. 1*D*). These data are consistent with the idea that WDR-48 decreases ubiquitination of USP-46 and promotes USP-46 protein levels independent of its own catalytic activity. Although we do not know the precise mechanism by which WDR-48 prevents USP-46 ubiquitination, we propose that WDR-48 binding to USP-46 either prevents the recruitment of an E3 ligase or blocks the ability of an E3 ligase to ubiquitinate key lysine residues on the surface of USP-46. Intriguingly, the crystal structure of WDR-48 bound to USP-46 reveals the presence of two USP-46 surface lysine residues (Lys226 and Lys235) at the interface of USP-46 and WDR-48 (35). It will be interesting in the future to test if WDR-48 inhibits ubiquitination of USP-46 by blocking access of an E3 ligase to these surface lysines. Interestingly, one or both of these lysine residues are conserved in USP-1 (Lys503) and USP-12 (Lys230 and Lys239), suggesting that WDR-48 may also stabilize these closely related USPs via the same mechanism. The WDR proteins and USP-46/USP-12 are conserved across phylogeny from yeast to humans, where they regulate several important processes including endocytosis, cell polarity, signal transduction and mitochondrial biogenesis (11,25,43,44). These WDR protein homologs may also regulate the stability of USP-46 homologs in non-neuronal cells in other species. For example, overexpression of WDR-48, but not WDR-20, can increase protein levels of human USP-12 resulting in stabilization of androgen receptors and proliferation of prostate cancer cells (16). In the filamentous fungi *Aspergillus nidulans*, protein levels of the USP-46 homolog CreB, were dramatically increased after expression of the WDR protein CreC (45). Although the precise mechanism controlling DUB levels was not investigated in these studies, our work suggests that these WDR proteins may increase protein levels of their interacting DUB partners by preventing their ubiquitination and degradation in the proteasome. We propose that this mechanism of stabilizing USP-46 and its homologs is likely conserved across phylogeny and may be a widely used mechanism to stabilize DUBs given that over 35% of all USPs interact with WDR proteins (24).

WDR-48 regulates USP-46 related DUBs via multiple mechanisms and is thus emerging as a critical DUB regulatory protein. First, WDR-48, either alone or together with WDR-20, can bind and stimulate the catalytic activity of USP-1, USP-46 and USP-12 (28–31,36). Second, WDR-48 can function as a substrate adaptor. For example, WDR-48 can bind substrates such as FANCD2/FANCI heteromers via its SUMO-like domain (SLD) and recruit them to USP-1 (46). Similarly, the C-terminal region of WDR-48 which includes the SLD is sufficient to bind the USP-12 substrate PHLPP1 (17). Third, WDR-48 can regulate the subcellular localization of its interacting USP. Bun107 and Bun62, the yeast homologs of WDR-48 and WDR-20, respectively, can recruit the USP-46 homolog Ubp9, to specific locations in the cytoplasm (25). The ability of WDR-48 to relocalize USPs is underscored by a study showing that human papilloma viruses (HPVs) have evolved to manipulate WDR-48 in order to control the subcellular localization of their interacting USPs. Specifically, HPV protein E1 binds to WDR-48 and recruits USP-1, USP-12 and USP-46 to viral origins of replication in the nucleus (47). These studies, together with our data showing that WDR-48 promotes USP-46 stability, reveal that WDR-48 is a versatile regulator of this group of USPs.

Our data suggest that USP-46 is continuously degraded and that increased expression of WDR-48 promotes the stability of the DUB. We suspect that WDR-48 levels are kept at a relatively low level resulting in low levels of USP-46. We propose that increased expression of WDR-48 stabilizes USP-46 as a mechanism to control DUB availability. We previously showed that overexpression of USP-46 alone does not increase its ability to regulate GLR-1 (36), suggesting that *in vivo* there are other limiting factors required to regulate USP-46. Although our data show that expression of WDR-48 alone is sufficient to increase the stability of USP-46, co-expression of WDR-48 and WDR-20 are required to fully stimulate the catalytic activity of the DUB and modulate glutamatergic behavior (36). Together, these data suggest that increasing the expression of WDR-48 alone is not sufficient to promote USP-46 function. A signal that increases expression of WDR-48 together with another signal that controls WDR-20 expression could function as an “AND” gate to precisely control USP-46 function. Although the relevant signals that regulate WDR-48 expression *in vivo* are not yet known, we speculate that WDR-48 expression in neurons may be regulated during development or in response to changes in synaptic activity. Interestingly, another DUB that can also regulate mammalian AMPA-type glutamate receptors, USP8, has been shown to be regulated by synaptic activity. In this case, activation of NMDARs leads to rapid dephosphorylation and activation of the DUB (48). Further studies will be necessary to identify the upstream signals that control expression of both WDR-48 and WDR-20 in neurons to regulate USP-46 function.

USP-46 and USP-12 have been implicated in several cancers (including colon cancer, prostate cancer and glioblastoma) and in regulating both glutamatergic and GABAergic signaling in the nervous system. The fact that DUBs are proteases makes them an attractive drug target; however, blocking the catalytic cysteine residue present in the active sites of most DUBs and many other proteases, could lead to non-selective effects. Our study shows that WDR-48 stabilizes USP-46 and that interaction of WDR-48 with USP-46 is required for this effect. Designing drugs that disrupt the interaction of WDR-48 with USP-46 could be an effective and more specific strategy to inhibit USP-46 function. We propose that targeting the interaction interface between WDR proteins and their DUB partners may be a promising and more specific approach to destabilize DUBs and thus inhibit their function.

## Experimental Procedures

### Strains

The following strains were used for experiments described in this manuscript: N2 (Bristol) wild type, *pzIs40* (*Pnmr-1*∷USP-46∷GFP) II, *ljIs114* (*Pgpa-13*∷FLPase; *Psra-6*∷FTF∷ChR2∷YFP) X (gift from William Schafer), *pzIs37* (*Pglr-1*∷USP-46∷GFP) III, *pzEx386 (Pnmr-*1∷GFP), *usp-46* (*ok2232*) III, *pzIs25* (*Pglr-1*∷WDR-20; *Pglr-1*∷WDR-48) I, *pzEx230* (*Pglr-1*∷WDR-20), *pzEx231* (*Pglr-1*∷WDR-48), *pzEx378* (*Pglr-1*∷USP-46(C38A)∷GFP), *pzEx456* (*Pglr-1*∷Myc-WDR-48), *pzEx457* (*Pglr-1*∷Myc-WDR-48(W266A, R224E, K282D)) (also referred to as 3Xmut). All strains were maintained at 20 °C as described previously (49).

### Constructs, Transgenes and Germ-line Transformation

*pzIs25, pzEx230, pzEx231* and *pzEx378* were described previously (36). *Pnmr-1*∷USP-46∷GFP (FJ#129) was generated by replacing the *Pglr-1* promoter in *Pglr-1*∷USP-46∷GFP (FJ#109) with *Pnmr-1* (~1 kb) from pBM16 (50) using SphI and BamHI sites. *pzIs40* was created by injecting FJ#129 (50 ng/μl) with the coinjection marker *Pttx-3*∷GFP (50 ng/μl) followed by integration using a UV Stratalinker. *pzIs40* was backcrossed 3 times prior to imaging. *pzIs37* was created by injecting FJ#109 (50 ng/ul) with the coinjection marker *Pttx-3∷GFP* (50 ng/μl) followed by integration using a UV Stratalinker. *pzIs37* was backcrossed 3 times prior to imaging. *pzEx378* was created by injecting FJ#109 (50 ng/μl) with the coinjection marker *Pttx-3*∷dsRed (50 ng/μl). *pzEx386* was created by injecting pBM16 (50 ng/μl) with the coinjection marker *Pttx3*∷GFP (50ng/μl).

pMT3-FLAG-USP-46 (FJ#66), pMT3-HA-WDR-20 (FJ#94), pMT3-Myc-WDR-48 (FJ#96), have been previously described (36). The mammalian expression vector pMT3 was kindly provided by Dr. Larry Feig. Myc-WDR-48 (~2 kb) was subcloned into the *C. elegans* expression vector pV6 for expression under the *glr-1* promoter to create *Pglr-1*∷Myc-WDR-48 (FJ#130) by first cloning into pBlueScript using HindIII and KpnI sites and then into FJ#109 using BamHI and KpnI sites. *pzEx456* was created by injecting FJ#130 (50 ng/μl) with the coinjection marker *Pmyo-2*∷NLS-mCherry (50 ng/μl). pMT3-Myc-WDR-48(R224E,W266A, K282R) also referred to as Myc-WDR-48(3Xmut) (FJ#131) was generated by using Quikchange (Invitrogen) to generate point mutations in the parent plasmid pMT3-Myc-WDR-48 (FJ#96)(36). Myc-WDR-48(R224E,W266A, K282R) (~2 kb) was subcloned into the *C. elegans* expression vector pV6 for expression under the *glr-1* promoter to create *Pglr-1*∷Myc-WDR-48 (R224E,W266A, K282R)(FJ#132) as described for FJ#130. *pzEx457* was created by injecting FJ#132 (50 ng/μl) with the co-injection marker *Pmyo-2*∷NLS-mCherry (50 ng/μl).

### Imaging

Fluorescence imaging of USP-46∷GFP was performed as follows. Briefly, L4 larval-stage animals were immobilized using 30 mg/ml 2,3-butanedione monoxamine (Sigma-Aldrich), and the ventral nerve cord (VNC) was imaged in the anterior region of the animals just posterior to the RIG neuronal cell bodies or in the cell body of the VNC neuron PVC. 1 μm (total depth) Z-series stacks were collected using a Carl Zeiss Axioscope M1 microscope with a 100X Plan Apochromat (1.4 numerical aperture) objective equipped with GFP and Cy3.5 filters. Images were collected with an Orca-ER charge coupled device (CCD) camera (Hamamatsu) and MetaMorph (version 7.1) software (Molecular Devices). Maximum intensity projections of Z-series stacks were used for quantitative analyses of fluorescence. Exposure settings and gain were adjusted to fill the 12-bit dynamic range without saturation and were identical for all images. The intensity for each worm was determined by averaging the maximum intensity from three separate ROIs. For proteasome inhibitor pretreatment, animals were placed on NGM plates containing 50 μM Bortezomib for 6 hours prior to imaging (51).

### Cell culture and Transfections

HEK293T cells were cultured in Dulbecco’s modified Eagle’s medium (DMEM) supplemented with 10% fetal bovine serum (FBS) and 100 U/ml penicillin. Cells were maintained in a humidified, 5% CO_2_ atmosphere at 37°C.

For protein synthesis inhibition experiments, HEK293T cells were seeded into a 100 mm dish and transfected with 6 μg of FLAG-USP-46 using Polyethylenimine (PEI, PolySciences) for 16-18 hours. Cells were trypsinized and plated into a 6 well plate and reverse transfected with Myc-WDR-48 (3 μg DNA), and/or HA-WDR-20 (2 μg DNA) using Lipofectamine 2000 (Invitrogen). 24 hours post-transfection the cells were again split into 12-well plates at 100,000 cells per well for cycloheximide time course studies. Cells were lysed in mammalian cell lysis buffer (MCLB, 50 mM Tris (pH 7.5), 150 mM NaCl, 0.5% Nonidet P-40, HALT Inhibitors (Pierce)). Cells were incubated at 4°C for 10 min and then centrifuged at 14,000 rpm for 15 min at 4°C. The supernatant was removed, and the protein concentration was estimated using the Bradford method (Bio-Rad).

For immunoprecipitation studies, HEK293T cells in 6-well plates were transfected with FLAG-USP-46, Myc-WDR-48, Myc-WDR-48(R224E, W266A, K282D) or HA-WDR-20, with a total of 2 μg of DNA per well, using Lipofectamine 3000 and harvested and lysed at 24-36 hours post-transfection for immunoprecipitations. HEK293T cells stably expressing human FLAG/HA-USP46 was created by lentiviral transduction as described elsewhere (52). Stable clones were selected using 1 μg/ml puromycin.

### Immunoprecipitation and Immunoblotting

For immunoprecipitation experiments, HEK239T cells were washed once with PBS and lysed after 24 h with MCLB. Lysed cells were centrifuged at 14,000 rpm for 10 min at 4°C. Cleared lysate was incubated for 4–12 hours with anti-FLAG antibody-coupled magnetic beads (Sigma) in the presence of 10 μg/ml HALT protease inhibitors and phosphatase inhibitors (sodium fluoride 50 μg/ml, sodium orthovanadate 5 μg/ml). Immunoprecipitated complexes were washed four times with MCLB and resuspended in 2X SDS-PAGE sample buffer. Samples were subjected to SDS-PAGE on 10% acrylamide gels and subsequently transferred to a nitrocellulose membrane. Membranes were blocked in Tris-buffered saline with Tween (TBS-T) and 5% milk prior to incubation with various primary antibodies.

For denaturing immunoprecipitation studies, FLAG/HA USP46 293T cells were lysed in MCLB containing 1% SDS, and vortexed at room temperature for 10 min. Insoluble material was removed by centrifugation (14,000rpm, 5 min). Protein concentration in the supernatants was determined by BCA assay (Pierce). Equal amounts of protein from each experimental sample were taken and diluted 10 fold in MCLB to acquire a final SDS concentration of 0.1%. Lysates were incubated with anti-FLAG antibody–coupled magnetic beads overnight at 4 ºC and then washed 3 times with MCLB. Beads were suspended in 2X SDS sample buffer, resolved by SDS-PAGE and transferred to nitrocellulose membranes. Membranes were blocked in Tris-buffered saline with Tween (TBS-T) and 5% milk prior to incubation with various primary antibodies.

Whole worm lysates were obtained by placing 100 animals in 2X SDS sample buffer and boiling at 95°C for 5 min, vortexing for 1 min and boiling again at 95°C for 5 min. Samples were subjected to SDS-PAGE and immunoblotting.

The following antibodies were used for immunoblotting: mouse-anti-GFP (Covance), rabbit anti-tubulin (ab4074, Abcam), rabbit anti-FLAG (M2, Sigma), mouse anti-Myc (clone 9E10, Santa Cruz), rat anti-HA (Covance), mouse anti-PCNA (clone PC10, Santa Cruz), and mouse anti-GAPDH (clone 0411, Santa Cruz), and mouse anti-ubiquitin (clone P4D1, Santa Cruz).

### Behavioral assays

All behavioral assays were performed using at least 10 young adult hermaphrodites over at least 3 independent experiments and by an experimenter who was blinded to the genotypes of the animals being tested. Optogenetic activation of the ASH-dependent nose touch response was performed similarly to what has been described previously (53). Gentle touch to the nose of the worm (nose-touch) (39,40,54) or photostimulation of ASH expressing channelrhodopsin2 (ChR2)(53,55) result in locomotion reversals away from the stimulus. This nose-touch response is dependent on presynaptic glutamate and GLR-1 in postsynaptic interneurons (39,40,54). We previously showed that mutants with decreased levels of GLR-1 in the VNC exhibit defects in the nose-touch response (12). Animals expressing ChR2 specifically in ASH sensory neurons (*ljIs114, Pgpa-13*∷FLPase; *Psra6*∷FTF∷ChR2∷YFP)(56) were grown for one generation in the dark on NGM agar plates spotted with OP50 and the ChR2 co-factor all-*trans*-retinal (ATR, 100 μM). Animals were subsequently transferred to an NGM agar plate spotted with OP50 (without ATR) and illuminated with 1 s pulses of blue light (0.47 mW/mm2) from a mercury bulb filtered through a GFP excitation filter (480 nm) under 32x total magnification on a Leica MZ16F microscope. A locomotor reversal was scored as a positive response if the backward movement was greater than the distance from the nose to the terminal bulb of the pharynx observed during or immediately after blue light illumination.

## Acknowledgements

We would like to thank William Schafer and the *Caenorhabditis* Genetics Center for strains, and Larry Feig for reagents. We thank Colleen Kreiser for help with preliminary experiments, and Eric Luth, Bethany Rennich and Betty Ortiz for advice and critical comments on this manuscript.

## FOOTNOTES

Funding for this work was supported in part by a National Science Foundation Grant (IOS1353862 to P.J.), National Institutes of Health Grants (R56NS059953 to P.J., and R01GM127557 to M.R.) and the Synapse Neurobiology Training Grant (T32 NS061764 to M.H.), the Tufts Center for Neuroscience Research Grant (P30NS047243) and the NIH Office of Research Infrastructure Programs Grant (P40OD010440) to the *Caenorhabditis* Genetics Center.

The abbreviations used are: DUB, deubiquitinating enzyme; WDR, WD40-repeat; VNC, ventral nerve cord; IP, immunoprecipitation; USP, ubiquitin-specific protein; CHX, cycloheximide; BTZ, bortezomib; UAF1, USP1-associated factor-1

---

**Supplemental Figure 1.**
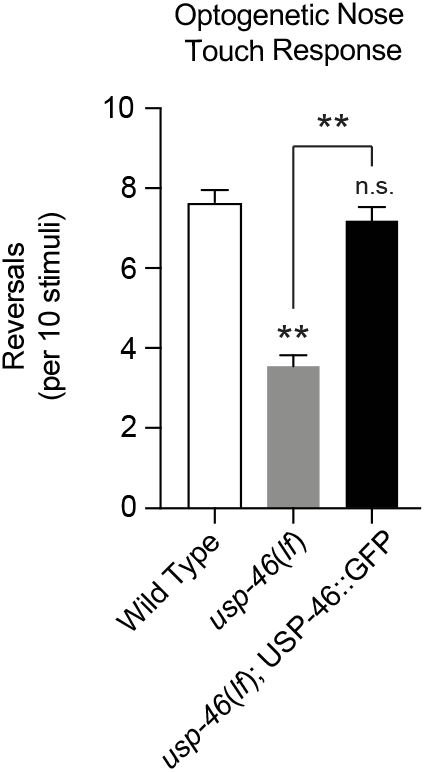
GFP-tagged USP-46 is functional in vivo. *A*, Optogenetic activation of the ASH-dependent nose touch response was performed, as described previously (53). Animals stably expressing channelrhodopsin2 (ChR2) in ASH sensory neurons (*ljIs114, Pgpa-13*∷FLPase; *Psra-6*∷FT-F∷ChR2∷YFP) were fed with OP50 along with the ChR2 co-factor all-*trans*-retinal (ATR, 100 μM). Single worms were stimulated with blue light and the number of reversals/10 stimuli was measured (see Experimental procedures). The average number of reversals per 10 stimuli are shown for the following genotypes: wild type (n=10), *usp-46*(*ok2232*, n=10), and USP-46∷GFP expressed under the *nmr-1* promoter (*pzIs40*);*usp-46*(*ok2232*, n=10). Values that differ significantly from the wild-type (ANOVA, Dunnett’s multiple comparison tests) are indicated as follows: **, *p* ≤ 0.001. n.s. p>0.05.

